# Phospho-enol pyruvate carboxykinase inhibition limits inflammatory T cell activation

**DOI:** 10.1101/2025.09.09.675120

**Authors:** Rebecca J. Brownlie, Helen Carrasco Hope, David Wright, Graham P. Cook, José C. Perales, Robert J. Salmond

**Author notes:** Correspondence: Robert J. Salmond. Contributed equally (Rebecca J. Brownlie, Helen Carrasco Hope).

## Abstract

Following antigenic stimulation, T cells switch from a catabolic metabolic state maintained by low levels of nutrient uptake to an anabolic metabolism that sustains the biosynthetic and energetic demands of clonal expansion, differentiation and effector function. Much progress has been made in understanding the transcriptional and enzymatic regulation of activated T cell metabolism. However less is understood of the role for regulators of anaplerosis and cataplerosis such as phospho-enol pyruvate carboxykinases (PEPCK) in T cells. In the current work, we show that mitochondrial isoform PEPCK-M is upregulated following T cell activation whilst cytosolic PEPCK-C is not expressed. PEPCK inhibitors limited CD8^+^ T cell cytotoxic capacity and both CD4^+^ and CD8^+^ T cell inflammatory cytokine production. Suppression of T cell effector functions by PEPCK inhibitors was associated with decreased maximal mitochondrial respiration. These data suggest that PEPCK-M acts as a metabolic rheostat to enable optimal T cell activation.

## Introduction

The processes of T cell activation and differentiation are linked to the regulation of cellular metabolism. These pathways provide the energy required for growth, proliferation and effector functions whilst metabolites can also directly modulate T cell differentiation. Importantly, dysregulation of cellular metabolism has been linked to immune decline in aging [1; 2], the failure of anti-tumour T cell responses [3; 4] and autoimmunity [5; 6].

The changes in T cell functional state that follow T cell antigen receptor (TCR) triggering are accompanied by metabolic reprogramming. TCR and costimulatory CD28 signals promote the activation of mTOR and Myc signaling pathways that result in upregulation of nutrient receptors and activation of glycolytic and glutaminolysis pathways [7; 8]. T cell metabolic reprogramming is also influenced by cytokines such as transforming factor factor *β* [9; 10]. Both CD4^+^ T helper (Th) cells and CD8^+^ cytotoxic T lymphocytes (CTLs) rely upon aerobic glycolysis to meet the metabolic demands of proliferation and effector differentiation whilst mitochondrial biogenesis and metabolism are also required for T cell activation [11; 12; 13; 14]. The tricarboxylic acid (TCA) cycle serves to link catabolism of carbohydrates, fats and proteins to oxidative phosphorylation (OXPHOS) via the production of NADH and FADH. Glycolysis and the TCA cycle also provide precursors for biosynthetic pathways. The removal of intermediate metabolites in metabolic pathways is termed cataplerosis, whilst effector T cells utilise these pathways concurrent with those that replenish these metabolites (anaplerosis). For example, T cell growth and proliferation is dependent upon anaplerotic production of the TCA cycle intermediate *α*-ketoglutarate from glutamine in glutaminolysis [15; 16].

A wealth of data has defined the role of transcriptional and enzymatic regulators of glycolysis and glutaminolysis in T cell activation. By contrast, the specific roles of regulators of anaplerosis and cataplerosis in T cells are poorly understood. Two isoforms of phosphoenolpyruvate carboxykinase (PEPCK) are central players in the regulation of this axis as enzymes that catalyse the conversion of oxaloacetate (OAA) to phosphoenolpyruvate (PEP) [17]. Cytoplasmic isoform PEPCK-C (encoded by *PCK1*/*Pck1*) is a critical enzyme in gluconeogenesis, whilst mitochondrial PEPCK-M (encoded by *PCK2*/*Pck2*) has a regulatory function in mitochondrial metabolism. In tumour cells, PEPCK-C promotes glutaminolysis and TCA cycle flux [18], whilst downregulation of TCA cycle anaplerosis/cataplerosis and a subsequent reduction in OXPHOS was associated with decreased PCK2 expression in melanoma [19]. Of note, ectopic overexpression of PEPCK-C can partially overcome the requirement for glycolysis in T cell activation. PEP produced during glycolysis or following enforced PEPCK-C expression facilitates prolonged TCR-induced Ca^2+^ signalling and nuclear factor of activated T cells (NFAT) activation [20]. High levels of PEP, produced during glycolysis or following PEP supplementation, have been shown to impede Th17 cell differentiation whereas a paucity of PEP, under conditions of glucose starvation, impairs Th1/CTL responses [20; 21], highlighting the key importance of regulating PEP levels in T cells. Furthermore, recent studies have shown that PEPCK-C expression is elevated in memory T cell populations where it functions to increase glycogen synthesis and metabolism to fuel the pentose phosphate pathway [22]. PEPCK-C expression is important in memory T cells for maintenance of high levels of glutathione (GSH) and redox homeostasis [22; 23]. By contrast, less is known of the role of PCK isoforms in effector and inflammatory T cell activation and metabolism.

In the current work, we assessed expression levels of PEPCK isoforms in mouse T cell subsets and used pharmacological inhibition to ascertain the importance of PEPCK function during T cell activation. Data show that PEPCK-M is the predominant isoform expressed in effector T cells whilst PEPCK inhibition limited CD8^+^ T cell cytotoxic capacity and both CD4^+^ and CD8^+^ T cell inflammatory cytokine production. Suppressive effects of PEPCK inhibitors on T cell function were distinct from previously described roles of PEPCK-C in maintaining GSH levels but were associated with decreased maximal mitochondrial respiration.

## Materials and Methods

### Database searches

Proteomic data were taken from the Immunological Proteome Resource (https://immpres.co.uk). Data shown are protein copy number/cell expressed by mouse T cell populations within haematopoietic cell proteomes dataset, using “Pck1” and “Pck2” as search terms. Data in figure 1B are PEPCK-M protein copy numbers within the Myc regulated T cell proteome dataset. RNA-Seq data shown in Figure 1C are taken from datasets originally published in Hope et al. 2022 [9] and represent fragments per kilobase million values for *Pck1* and *Pck2* transcripts within OT-I T cells stimulated for 24h with SIITFEKL peptide.

**Figure 1.**
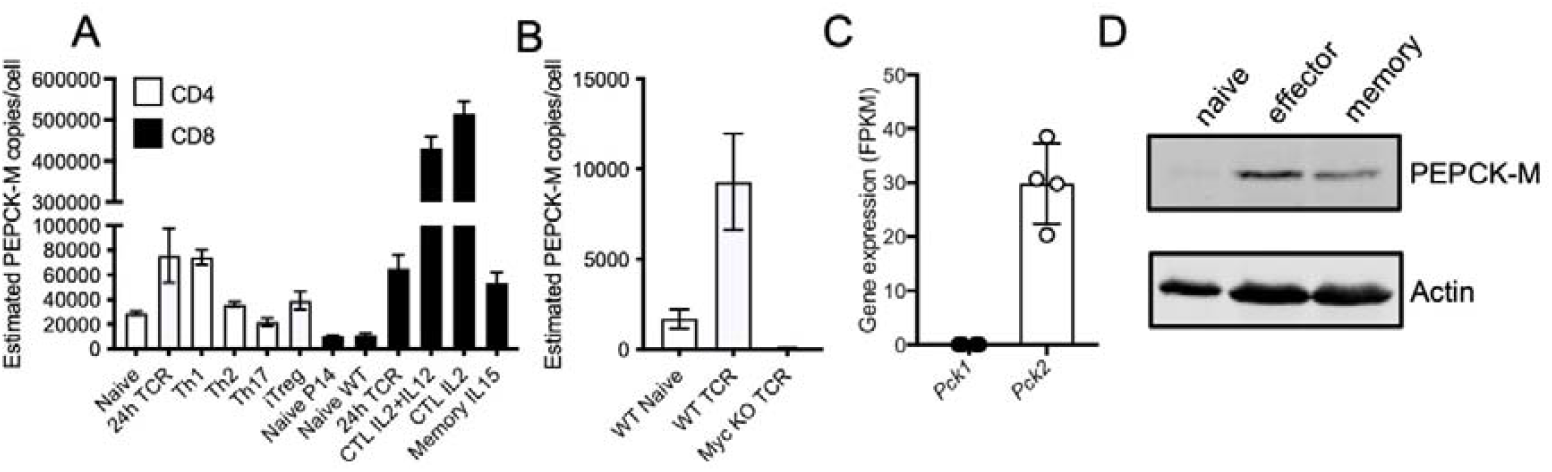
Expression of PEPCK-M in effector T cells. PEPCK-M protein copy numbers / cell in mouse T cell subsets **(A)** and in control and Myc-deficient CD8^+^ T cells stimulated for 24h with CD3 and CD28 antibodies **(B).** Data are from Immunological Proteome Resource (immpres.co.uk). Values are means and error bars represent SD from n=3 biological replicates. **(C)** Expression of *Pck1* and *Pck2* transcripts expressed as fragments per kilobase million (FPKM) in OT-I T cells activated for 24h with SIITFEKL antigen. Circles represent individual values from 4 biological replicate samples. Data are from RNA-Sequencing datasets originally published in. **(D)** Western blot analysis of PEPCK-M expression in naïve and day 6 effector or memory-phenotype CTLs. *β*-actin reprobes serve as protein loading controls.

### Mice

OT-I *Rag1*^*-/-*^ and C57BL/6J mice were maintained at the University of Leeds’ St. James’s Biomedical Services animal facility. All experiments were performed using both male and female mice at 7-15 weeks of age.

### Cell Lines

ID8-OVA-fLuc cells [24] were maintained in Iscove’s Modified Dulbecco’s Medium (IMDM, Gibco) supplemented with 5% foetal calf serum (FCS) (Gibco), L-glutamine and antibiotics (100U penicillin, 100 *μ*g/ml streptomycin). EL4-OVA cells, originally from Klaus Okkenhaug (University of Cambridge), were maintained as above with the addition of 400 *μ*g/mL G418 antibiotic. Cell lines were negative for mouse pathogens, including mycoplasma contamination.

### T cell culture and stimulation

OT-I T cells were obtained from lymph nodes of OT-I Rag1^-/-^ mice. Single cell suspensions were prepared by dissociating lymph nodes using a 70 *μ*M filter, with cell purities of CD8^+^ T cells being >90%. OT-I T cells were cultured in IMDM supplemented with 5% FCS, L-glutamine, antibiotics (100U penicillin, 100 *μ*g/ml streptomycin) and 50 *μ*M 2-mercaptoethanol. OT-I T cells were activated using SIINFEKL peptide (Cambridge Peptides) for time periods indicated in Figure legends. In some experiments, control OT-I T cells were left unstimulated *in vitro*. 3-mercaptopicolinic acid (Cayman Chemicals) was added to culture media at a final concentration of 100 *μ*M unless otherwise stated. PEPCK inhibitor 5-chloro-*N*-{4-[(cyclopropylmethyl)-1-(2-fluorbenyzl)-2,6-dioxo-2,3,6,9-tetrahydro-1*H*- purin=8=yl)methyl]phenyl}-1,3-dimethyl-1*H*-pyrazole-4-sulfonamide (iPCK2) was synthesized as previously described [25] and was added to culture media at a final concentration of 5 *μ*M. Where indicated, 1 *μ*M L-glutathione (GSH, Tocris), 50 *μ*M glycogen phosphorylase inhibitor (Cayman Chemicals) or 1mM phospho-enol pyruvate monopotassium salt (PEP, Merck) were added to culture. For generation of cytotoxic T lymphocytes (CTLs), OT-I T cells were activated ± 3-MP or iPCK2 with 10^-8^M SIINFEKL for 2 d, followed by 4d differentiation with 20 ng/mL recombinant human IL-2. For analysis of cytokine production, day 6 CTLs were re-stimulated with 10^-7^M SIINFEKL for 4h in the presence of 2.5 *μ*g/mL brefeldin A (Sigma). For Th1 and Th17 differentiation, CD4^+^ T cells were purified from C57BL/6 lymph nodes by negative selection using magnetic beads (Miltenyi Biotech). T cells were activated in 48 well plates for 3d (Th1) or 6d (Th17) with platebound CD3*ε* (Clone 145-2C11, BioLegend) and CD28 mAb (Clone 37.51, BioLegend) with the following recombinant cytokines (all Peprotech): Th1 – 10 ng/mL mouse IL-12, 10 ng/mL human IL-2; Th17 – 1 ng/mL mouse TGF*β*, 20 ng/mL mouse IL-6, 5 *μ*g/mL anti-IFN*γ* (Clone XMG1.2, BioLegend), 5 *μ*g/mL blocking anti-CD25 (Clone PC61, BioLegend). Brefeldin A was added to Th1 and Th17 cultures for the final 4hours of culture prior to intracellular staining and flow cytometry.

### Western blotting

Lysates were prepared in RIPA lysis buffer, protein concentrations assessed by Bradford Assay (Thermo Fisher) and 15 *μ*g protein / sample loaded on polyacrylamide gels. Western blotting was performed as previously described, using a Li-COR Odyssey Imaging System [26]. Antibodies used were: rabbit polyclonal anti-PECK-M/PCK2 (#6924, Cell Signaling Technology); mouse anti-*β*-actin (Clone AC-15, Sigma), goat anti-rabbit AF680, goat anti-mouse AF790 (Molecular Probes).

### Flow cytometry

The following antibodies were used: CD4-allophycocyanin (APC) (Clone GK1.5), CD8*β*- phycoerythrin (PE) cyanine 7 (Cy7) (Clone YTS156.7.7), CD69-peridinin chlorophyll protein (PerCP) Cy5.5 (Clone H1.2F3), CD71-fluorescein isothiocyanate (FITC) (Clone R17217), PD-1-PE (29F.1A12), granzyme B-Pacific Blue (Clone GB11), Tbet-PE (Clone 4B10), IFN*γ*-alexa flour 488 (AF488) (Clone XMG1.2), TNF-PerCP Cy5.5 (Clone MP6-XT22), IL-17A-APC (Clone TC11-18H10.1) (all BioLegend) and retinoic acid-related orphan receptor (ROR) gamma-PE (Clone B20, eBioscience). For live cell discrimination, cells were stained with live/dead aqua dyes (Life Technologies). For intracellular staining cells were fixed in eBioscience FoxP3 fix/permeabilisation buffers prior to staining in permeabilisation buffer. Samples were acquired with an LSRII (Becton Dickinson) or Cytoflex S (Beckman) flow cytometers and data were analysed using FlowJo Software (Treestar).

### IL-2 ELISA

OT-I T cells were activated with 10^-6^M SIINFEKL ± 3-MP in the presence of 2.5 *μ*g/mL CD25 blocking antibody, to prevent IL-2 consumption, and supernatants collected at 24h. Levels of supernatant IL-2 were assessed by ELISA using the mouse IL-2 DuoSet kit, according to manufacturers’ instructions (R&D Systems).

### Quantitative RT-PCR

OT-I T cells were activated with 10^-6^M SIINFEKL ± 3-MP for 24h and cell pellets stored at - 80°C. RNA was prepared using the PureLink® RNA Mini Kit (Invitrogen) and cDNA synthesis was performed using the RevertAid First Strand cDNA synthesis kit (Thermo Scientific). *Gzmb* expression was quantified using the ddCT method and Taqman reagents using the QuantStudio 7 Real-Time PCR system ((Applied Biosystems). Relative expression of *Rpl13a* was used to normalize gene expression across samples. The following Taqman probes were used: *Gzmb* – Mm00442837_m1; *Rpl13a* – Mm05910660_g1.

### In vitro cytotoxicity assay

Target ID8-OVA-fLuc cells were seeded in 48 well plates for 6h, prior to addition of day 6 CTLs at the ratios indicated in figures. Following overnight culture, plates were gently washed in PBS to remove T cells and target cell debris, prior to assessment of luciferase activity by addition of luciferin (Regis Technologies) and IVIS imaging. Specific cell lysis was calculated by comparison of luminescence wells in experimental wells to target only and blank wells.

### EL4-OVA tumour model and ACT experiments

EL4-OVA cells (1×10^6^) were injected subcutaneously into the flank of C57BL/6 mice. After 5 days, mice were randomly divided into 3 groups; control mice received no ACT, whilst additional groups received intravenous injections via the tail vein of either control or 3-MP CTLs (5×10^6^/mouse). Tumour mass was assessed by caliper measurements every 2-3 days, until tumours in any control group mice reached a diameter of 15mm.

### Seahorse metabolic assays

Mitostress test kits (Agilent) were used to measure metabolic profiles using a Seahorse XFe96 analyser. Activated T cells were washed (3x) in PBS, prior to transfer to Seahorse assay plates (1×10^5^/well) and adhered using Cell-Tak solution (22.4 *μ*g/mL, Corning) in complete XF assay medium. Oligomycin (1 *μ*M), Carbonyl cyanide-p-trifluoromethoxyphenylhydrazone FCCP (1.5 *μ*M) and rotenone/antimycin A (500 nM) were injected using the Mitostress test protocol. Data were collected in Wave software and analysed using GraphPad Prism.

### Statistics

Statistical significance (p-value < 0.05) was determined by Student’s t-test, one-or two-way ANOVA with Tukey’s multiple comparisons tests using GraphPad Prism. Dots in graphs represent replicate samples and error bars represent SDs, unless otherwise stated.

### Study Approval

Mouse breeding and experiments performed were reviewed and approved by the University of Leeds Animal Welfare and Ethical Review Committee and were subject to the conditions of UK Home Office Project Licence PDAD2D507, held by RJS.

## Results

### Expression of PEPCK-M, but not PEPCK-C, in effector T cells

We used publicly available datasets to assess expression of cytosolic and mitochondrial PEPCK isoforms in mouse T cells. Quantitative mass spectrometry data from the Immunological Proteome Resource (ImmPRes) [27] showed that mitochondrial PEPCK-M was expressed at low levels in naïve CD4^+^ and CD8^+^ T cells (Fig. 1A). Expression levels of PEPCK-M were elevated in a majority of activated mouse T cell populations, as compared to naïve cells, with the highest levels being in effector CD8^+^ cytotoxic T lymphocyte (CTL) populations. By contrast PEPCK-C was not detected in any of the T cell mass spectrometry datasets available. Of note, analysis of a further proteomic dataset, originally published by Marchingo and colleagues [28] and available in ImmPRes, showed that upregulation of PEPCK-M in CD8^+^ T cells following TCR stimulation was strictly Myc-dependent (Fig. 1B). Consistent with the proteomic data, analysis of our published RNA-sequencing data [9] demonstrated that *Pck1* transcripts were absent whereas *Pck2* transcripts were readily detectable in activated CD8^+^ OT-I TCR transgenic T cells (Fig. 1C). Furthermore, western blot analysis demonstrated that PEPCK-M protein was expressed at low but detectable levels in naïve OT-I T cells and upregulated in effector and memory-phenotype CTLs (Fig. 1D). Together, these data indicate that mitochondrial PEPCK-M is the predominant PEPCK isoform expressed in effector T cells and that PEPCK-M/*Pck2* upregulation is part of the TCR-driven, Myc-dependent transcriptional programme that regulates T cell metabolic reprogramming.

### PEPCK inhibitors limit CD8 T cell cytolytic activity

To assess the role of PEPCKs in T cell activation we used the well characterized inhibitor 3-mercaptopicolinic acid (3-MP) [29; 30]. 3-MP acts as a competitive inhibitor of PEP/OAA binding and binds a second allosteric site in the PEPCK structure [29]. CD8^+^ OT-I T cells were stimulated *in vitro* with cognate SIINFEKL peptide ± 3-MP for 48 hours and T cell activation assessed. Flow cytometry analysis demonstrated that 3-MP did not impact on T cell growth, as assessed by FSC-A analysis, nor upon TCR-induced upregulation of activation marker CD69, transferrin receptor CD71, exhaustion-associated immune checkpoint receptor PD-1 or transcription factor Tbet (Fig. 2A). Furthermore, 3-MP did not affect TCR-induced OT-I T cell IL-2 production (Fig. 2B). By contrast, 3-MP limited TCR-induced upregulation of cytolytic effector granzyme B protein (Fig. 2C) and mRNA (Fig. 2D) expression, in a dose-dependent manner (Fig. 2E). Importantly, a second structurally distinct PEPCK inhibitor, iPCK2 [25], also selectively inhibited TCR-induced granzyme B but not PD-1 expression in OT-I T cells (Fig. 2F, 2G).

**Figure 2.**
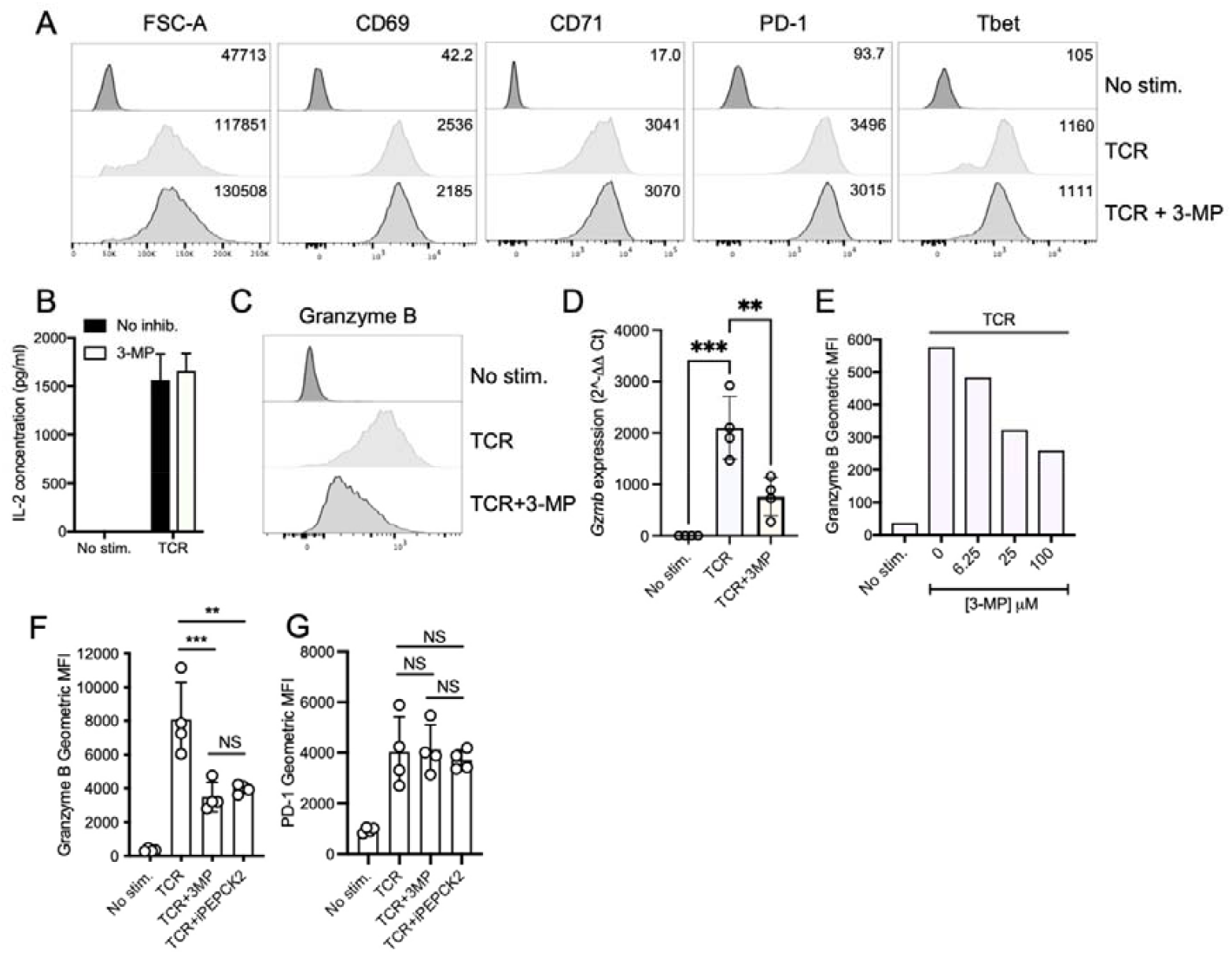
PEPCK inhibitors reduce T cell granzyme B expression. OT-I TCR transgenic T cells were stimulated with cognate peptide SIINFEKL (TCR) ± 3-MP or iPCK2 for 24h **(B)** or 48h (**A, C-F**). (**A**) Data shown are representative histograms (from n>8 experiments) of activated or control OT-I T cells’ forward scatter area (FSC-A) or expression of the indicated cell surface or intracellular markers, as assessed by flow cytometry analysis. Values in histograms are mean fluorescence intensities. (**B**) Levels of supernatant IL-2 were assessed by ELISA. Error bars represent SD (n=4 technical replicates from 1 of 2 repeated experiments). (**C**) Representative histograms from n>8 experiments of intracellular granzyme B. Values in histograms are mean fluorescence intensities. (**D**) 3-MP limits TCR-induced Gzmb transcription as determined by qRT-PCR. (**E**) Dose-dependent inhibition of granzyme B expression by 3-MP. iPCK2 and 3-MP TCR-induced impede granzyme B expression (**F**) but not PD-1 (G). Dots represent biological replicate values (D) or technical replicate values from 1 of 3 repeated experiments (F, G). NS – not significant, ** p<0.01, *** p<0.001.

To determine if this reduced level of granzyme B expression impeded cytotoxic activity, effector OT-I CTLs were generated in the presence or absence of 3-MP by stimulating cells for 2d with SIINFEKL peptide followed by 4d of differentiation using high dose IL-2, as per our previously described protocol [31]. As was the case following 48h of activation, day 6 CTLs generated in the presence of 3-MP (hereafter termed “3-MP CTLs”) had reduced levels of granzyme B as compared to control CTLs (Fig. 3A). Furthermore, 3-MP OT-I CTLs had impaired capacity to kill OVA-expressing ID8 tumour cell targets *in vitro*, as shown by a shift in the titration curve (Fig. 3B).

**Figure 3.**
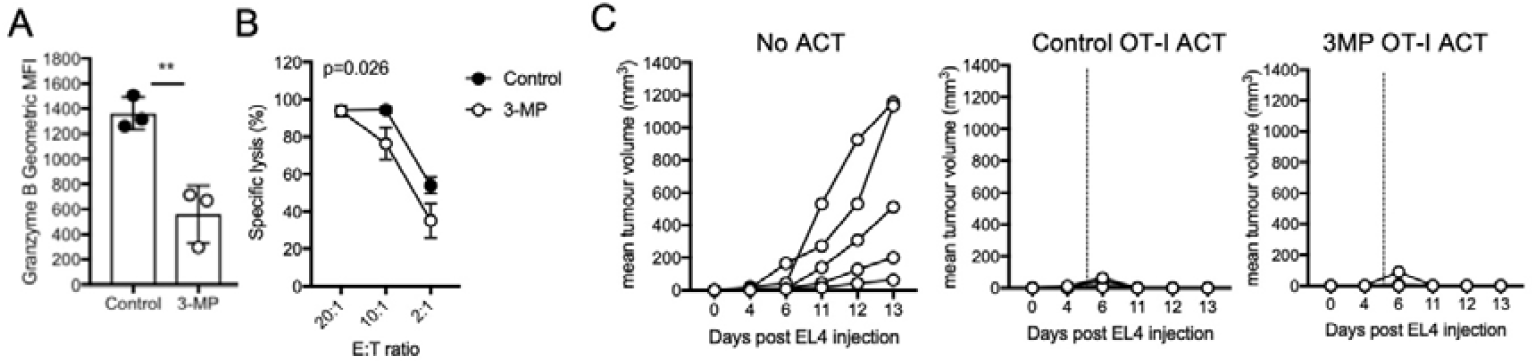
PEPCK inhibition reduces CD8 T cell cytolytic capacity. (**A**) Effector CTLs were generated by 6day stimulation in the presence or absence of 3-MP. Circles represent individual values from technical replicates of granzyme B expression (n=3 technical replicates, from 1 of >4 repeated experiments). ** p<0.01 by Student’s t-test. (**B**) Specific lysis of ID8-OVA target cells by control or 3-MP OT-I CTLs at the stated effector:target ratios (E:T). Statistical analysis was by 2-way ANOVA. (**C**) Groups of mice (n=5/group) were challenged with s.c. EL4-OVA and following 5 days received no ACT, control OT-I or 3-MP OT-I CTL ACT. Data shown are mean tumour volume as assessed by caliper measurements.

Next, we assessed the capacity of 3-MP CTLs to clear tumours *in vivo* using an adoptive T cell transfer (ACT) model. EL4-OVA lymphoma cells were injected into the flanks of C57BL/6 mice and allowed to establish tumours for 5d, following which mice received intravenous ACT using high numbers (5×10^6^) of either control or 3-MP OT-I CTLs. Data indicated that both control and 3-MP CTLs were competent to control EL4-OVA tumour growth in all mice assessed (Fig. 3C). Taken together these data indicate that 3-MP has a selective inhibitory effect on CD8^+^ T cell activation, reducing granzyme B expression and limiting CTL activity. However, at high effector:target ratios or following ACT using high numbers of CTLs, 3-MP CTLs retain the capacity to kill cancer cells both *in vitro* and *in vivo*.

### PEPCK inhibition limits T cell inflammatory cytokine production

We set out to determine whether PEPCK inhibition impacted upon acquisition of other T cell effector functions. To this end, day 6 OT-I CTLs generated in the presence or absence of 3-MP were re-stimulated with SIINFEKL peptide and effector cytokine production assessed by intracellular staining and flow cytometry. Whilst the proportions of 3-MP CTLs competent to produce IFN*γ* were only slightly reduced, the levels of IFN*γ* produced on a per cell basis, as assessed by mean fluorescence intensity of staining, were reduced by ∼50% as compared to control CTLs (Fig. 4A-C). By contrast, similar proportions of 3-MP and control CTLs produced TNF following re-stimulation, whilst the per cell level was mildly reduced in 3-MP treated cells as compared to controls (Fig. 4A-C). Further experiments demonstrated that CTLs grown in the presence of an alternative inhibitor, iPCK2, also had a reduced capacity to produce IFN*γ* upon antigenic restimulation, as compared to control CTLs (Fig. 4D).

**Figure 4.**
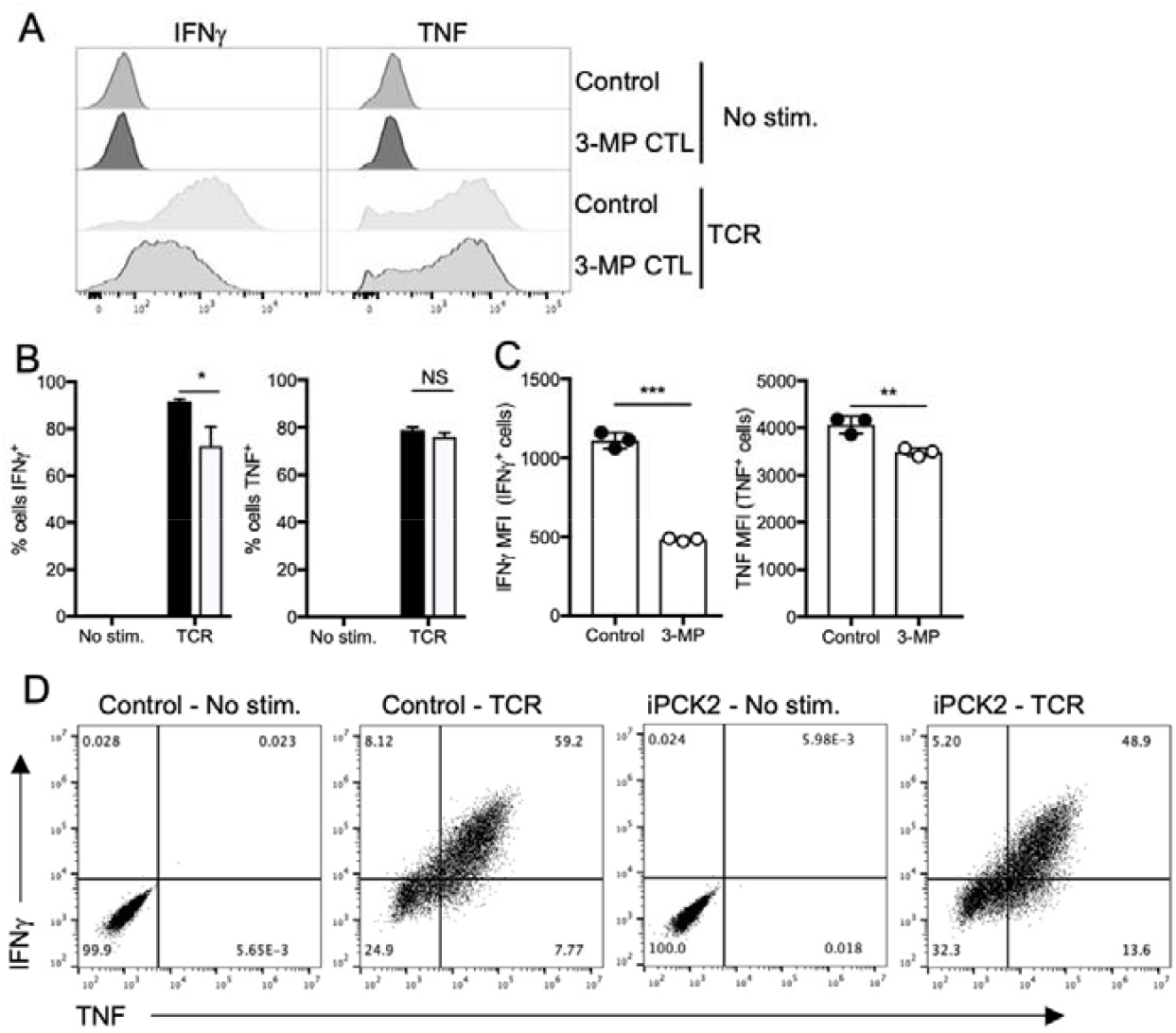
PEPCK inhibition limits CD8^+^ T cell inflammatory cytokine production. Effector CTLs were generated by 2 days of stimulation with SIINFEKL peptide followed by 4d of differentiation and expansion with IL-2 in the presence or absence of 3-MP (**A-C**) or iPck2 (**D**). Cell were restimulated with SIINFEKL (TCR) and levels of IFN*γ* and TNF production assessed by flow cytometry. (**A**) Representative histograms of IFN*γ* and TNF expression of control or 3-MP CTLs. Graphs show proportions of IFN*γ*^+^ and TNF^+^ OT-I control or 3-MP CTLs (**B**) and mean fluorescence intensity of cytokine staining in gated positive cells (**C**). Dots are representative of technical replicates and bars represent means ± SD. (**D**) Representative dotplots of IFN*γ* and TNF expression in control or iPck2-CTLs. Data are representative of 4 (**A-C**) and 2 (**D**) repeated experiments. NS – not significant; * p<0.05, ** p<0.01, *** p<0.001 as determined by Student’s t-test.

It was of interest to determine whether 3-MP had similar effects on CD4^+^ T cell cytokine production. Polyclonal C57BL/6 lymph node CD4^+^ T cells were activated with CD3 and CD28 antibodies in the presence of IL-12 for 3d and levels of the canonical Th1 cytokine IFN*γ* and transcription factor Tbet assessed by flow cytometry. As with the results determined for CD8^+^ T cells, 3-MP did not impact upon the upregulation of Tbet expression in CD4^+^ T cells but substantially impeded IFN*γ* production (Fig. 5A, B). In parallel experiments, the impact of 3-MP on Th17 cell differentiation was assessed. CD4^+^ T cells were activated under Th17 polarising conditions for 6d and levels of key Th17-associated transcription factor ROR*γ*t and IL-17A assessed by flow cytometry. Similar to the results for Th1 differentiation, levels of ROR*γ*t were comparable in control and 3-MP Th17 cells whereas the proportion of IL-17A^+^ cells was reduced by ∼50% by 3-MP (Fig. 5C, D). Together these results demonstrate that PEPCK inhibition does not impede the upregulation of lineage-defining transcription factors such as Tbet or ROR*γ*t during T cell differentiation, but instead reduces the capacity of effector CD8^+^ and CD4^+^ T cells to produce inflammatory cytokines.

**Figure 5.**
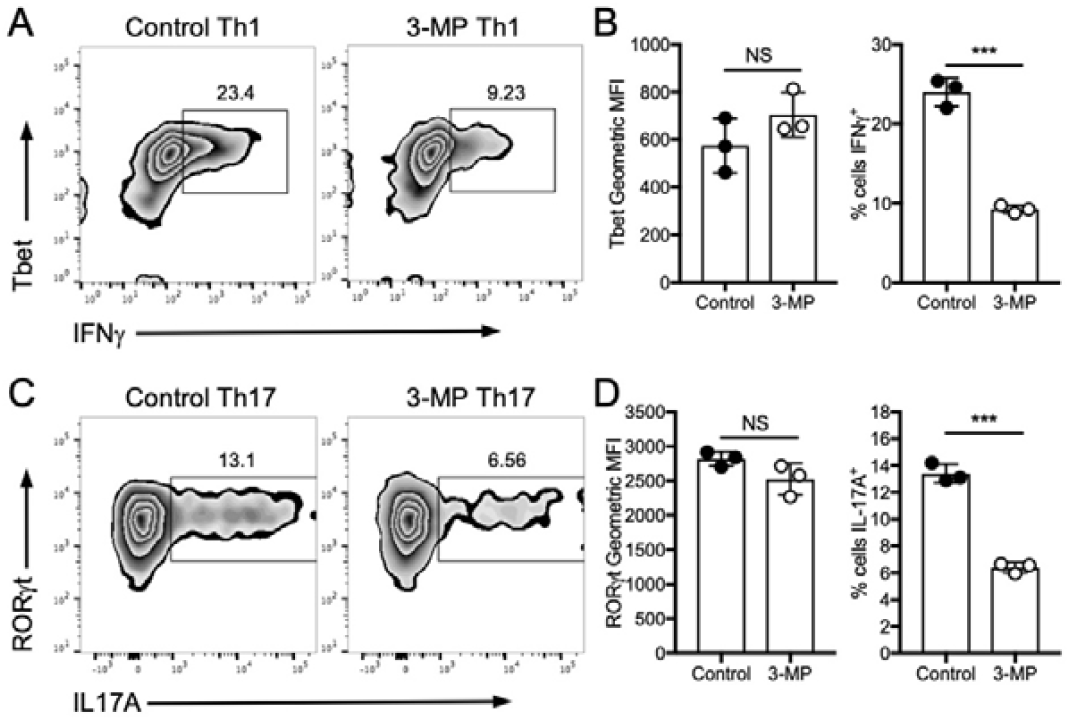
PCK inhibition limits CD4^+^ T cell inflammatory cytokine production. Purified C57BL/6 CD4^+^ T cells were stimulated under Th1 polarizing conditions for 3d (**A, B**) or Th17 polarizing conditions for 6d (**C, D**) and levels of lineage-specific transcription factor and cytokines assessed by flow cytometry. In all graphs, circles represent individual replicate values. Data are representative of 4 (**A-B**) and 3 (**C**,**D**) repeated experiments. NS – not significant; * p<0.05, ** p<0.01, *** p<0.001 as determined by Student’s t-test.

### Metabolic effects underpin inhibitory effects of 3-MP

We sought to define mechanisms underpinning the effects of 3-MP on inflammatory T cell effector responses. Previous reports indicated that the effects of PEPCK-C inhibition on memory T cells could be reversed by the addition of GSH to T cell cultures and were phenocopied by glycogen phosphorylase inhibitors (GPI) [22]. Therefore, we assessed the impact of GSH supplementation and GPI on TCR-induced OT-I T cell granzyme B expression in the presence or absence of 3-MP. The addition of GSH did not reverse 3-MP-mediated inhibition, whilst GPI had no effect on TCR-induced granzyme B expression (Fig. 6A). These data indicate that the effects of 3-MP reported here are mechanistically distinct from the effects of PEPCK-C deletion in previous studies. A further possibility was that 3-MP treatment limited T cell activation by impacting on the cellular abundance of PEP. However, the addition of PEP to T cell culture media did not alleviate the effects of 3-MP inhibition and, when given alone, had an inhibitory effect on TCR-induced granzyme B expression (Fig. 6B).

**Figure 6.**
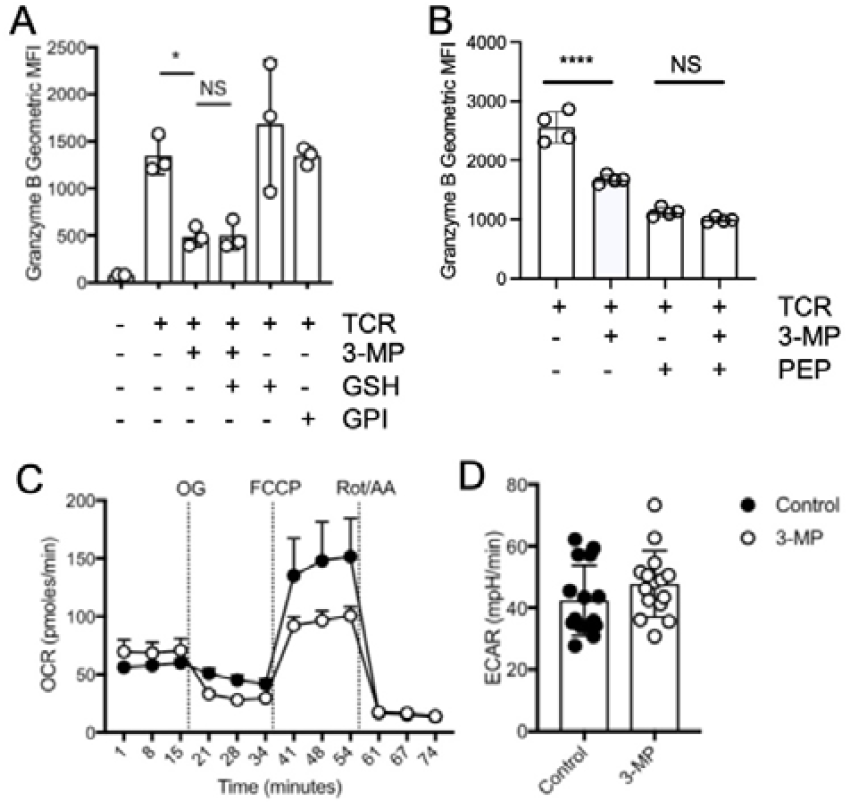
3-MP impacts upon mitochondrial respiration. OT-I T cells were activated with SIINFEKL peptide (TCR) in the presence or absence of 3-MP, GSH and glycogen phosphorylase inhibitor (GPI) (**A**) or 3-MP and PEP (**B**) for 48h and granzyme B assessed by intracellular staining and flow cytometry. Circles represent individual replicates values from 1 of 3 repeated experiments. NS – not significant, * p<0.05 as assessed by T-test, with Holm-Sidak correction for multiple comparisons (**C**) OT-I T cells were activated for 24h ± 3-MP then analysis of oxygen consumption rates (OCR) was performed using the Seahorse Mitostress test. Values represent mean and error bars represent SEM of technical replicates (n=5) from one of 3 repeat experiments. Baseline extracellular acidification rates (ECAR) in the presence of glucose were also assessed, at time-points prior to addition of oligomycin (OG) (**D**).

Metabolic analyses using a Seahorse analyser indicated that OT-I CD8^+^ T cells activated for 24h in the presence of 3-MP had a reduced maximal oxygen consumption rate (OCR), an indicator of OXPHOS activity, relative to control cells (Fig. 6C). By contrast, control and 3-MP treated OT-I T cells had similar levels of basal glycolytic activity as assessed by Seahorse analysis of extracellular acidification rate (ECAR) (Fig. 6D). These data suggest that a primary function for PEPCKs in activated T cells is to act as a metabolic rheostat in the face of energetic challenge, whilst 3-MP treatment impedes these functions.

## Discussion

In this paper, we have shown that PEPCK-M is the primary PEPCK isoform expressed in effector T cell populations, whilst PEPCK inhibition selectively impedes T cell cytotoxic function and inflammatory cytokine production. These data implicate PEPCKs as key metabolic regulators of T cell effector function and as potential targets to modulate T cell metabolism and function.

PEPCK-C is a rate-limiting enzyme in hepatic gluconeogenesis with high protein expression detected in kidney, gut and liver [32]. A previous study indicated that this cytosolic isoform was upregulated in memory T cells but was low/undetectable in naïve and effector T cells [22]. Here, searches of publicly available and published proteomic and RNA-Seq datasets determined that PEPCK-C protein and *Pck1* mRNA were not detected in naïve and effector mouse T cell subsets, whereas PEPCK-M / *Pck2* was widely expressed. These data suggest that PEPCK-C has an important role in T cell memory, whereas PEPCK-M may have a more general role in T cell effector functions. Consistent with this, *Pck1*-haploinsufficiency does not impair effector T cell responses during bacterial infections but limits the generation of memory T cell populations [22].

3-MP was established as an inhibitor of PEPCKs in the 1970s [30; 33] and, more recently, was shown to act as a competitive inhibitor of PEP/OAA binding, and to bind a second allosteric site in the PEPCK structure [29]. In the present work, we showed that 3-MP did not have a global inhibitory effect on T cell activation but instead had a selective effect on expression of granzyme B in CD8^+^ T cells and inflammatory cytokines by both CD4^+^ and CD8^+^ T cells. A second structurally distinct inhibitor, iPCK2 [25], also selectively reduced granzyme B expression and cytokine production, strengthening the evidence for a role for PEPCKs in these processes. CTLs generated in the presence of 3-MP were less cytotoxic than control counterparts on a cell:cell basis but were able to kill target cells *in vitro* and control tumour growth *in vivo* if supplied in sufficient numbers. Furthermore, 3-MP did not impair Th1 and Th17 differentiation *per se*, as levels of lineage-defining transcription factors Tbet and ROR*γ*t were not impaired, but rather reduced the capacity of T cells to produce high levels of cytokines. Previous studies linked PEPCK-C expression in memory T cells to the maintenance of GSH levels and redox balance via the regulation of glycogen synthesis and metabolism [22]. By contrast, neither GSH nor glycogen phosphorylase inhibition affected granzyme B expression in control or 3-MP treated T cells, suggesting a different mechanism being key to the results of the present study. PEP supplementation impeded TCR-induced granzyme B expression suggesting that 3-MP does not inhibit T cell activation via limiting PEP levels. Indeed, given that 3-MP did not appear to impede glycolytic flux, it is likely that enolase-dependent PEP production compensates for any direct effect on PEP levels resulting from PCK inhibition. Of note, cytotoxic effector proteins such as granzyme B are amongst the most abundant proteins expressed by CTLs [34]. Selective reductions in granzyme B and cytokine expression following 3-MP treatment implies that PCKs function as rheostats in the face of energetic challenges such as when the demand for gene transcription and protein synthesis is high. Consistent with this, our metabolic analyses determined that 3-MP-treated CD8^+^ T cells had reduced maximal, stressed mitochondrial respiration rates. Similarly, previous studies determined that downregulation of PEPCK-M impairs OXPHOS in cancer cells [19].

Our working hypothesis is that inhibition of PEPCK-M underlies the effects of 3-MP and iPCK2 on T cell activation. This is based on the observation that PEPCK-M but not PEPCK-C is expressed abundantly in effector T cells. Nonetheless we cannot formally rule out a role for either isoform at present. Recent studies have shown that *Pck2*^*-/-*^ mice are viable and fertile [35], and that PEPCK-M plays a role in maintenance of a mitochondrial PEP cycle, alongside pyruvate carboxylase and pyruvate kinase, that is important for regulating pancreatic *β* cell insulin secretion [35; 36]. A role for PEPCK-M in regulating LPS-driven Kupffer cell inflammatory responses has been suggested [37]. Future work will require in depth analysis of immune phenotypes of *Pck2*^*-/-*^ mouse strains to address the question of the precise role of PEPCK-M in adaptive immune responses.

Whilst 3-MP treated OT-I T cells had impaired effector functions *in vitro*, our adoptive T cell experiments showed that they were still competent to clear EL4-OVA tumours. It is possible that *in vivo* defects of 3-MP CTLs might be revealed by using lower numbers of CTLs in ACT experiments, or by treating tumour-bearing mice directly with PEPCK inhibitors. In this regard, several recent studies have reported differing effects of 3-MP treatment on *in vivo* T cell responses to tumours. Ma and colleagues reported that 3-MP treatment resulted in accelerated growth of B16-OVA melanoma, whilst Pck1 heterozygous OT-I T cells were inferior to control cells in controlling tumour growth following ACT [22]. Adoptive transfer of PEPCK-C-overexpressing tumour-reactive T cells has been shown to result in more effective tumour control than control T cell ACT in several studies [20; 22]. By contrast, a recent study reported that 3-MP treatment decreased Treg proliferation and increase the proportions of IFN*γ*-producing CD8^+^ T cells within B16 tumours, without impacting overtly on overall tumour growth [38]. Interpretation of these studies is complicated as 3-MP will have direct and indirect effects on many different cell populations *in vivo*, and the overall outcome on tumour growth and anti-tumour immunity will be highly context-dependent. Nonetheless, these previous studies and the work presented here add to a growing body of evidence that implicates PEPCKs as key regulators of T cell metabolism and activation that can be targeted to influence T cell responses *in vivo*.

## Data availability statement

## Ethics Statement

The animal study was reviewed and approved by the University of Leeds AWERB.

## Author Contributions

RJS designed the study. RJB, HCH, DW and RJS acquired and analysed the data. GPC and RJS supervised the work. JCP provided essential reagents and expert advice in inhibitor studies. RJS wrote the first draft of the manuscript. All authors read and discussed manuscript drafts. All authors contributed to the article and approved the submitted version.

